# *Streptococcus pyogenes* ϕ1207.3 is a temperate bacteriophage carrying the macrolide efflux gene pair *mef*(A)-*msr*(D) and capable to lysogenise different Streptococci

**DOI:** 10.1101/2022.10.13.512196

**Authors:** Francesco Santoro, Gabiria Pastore, Valeria Fox, Marie-Agnes Petit, Francesco Iannelli, Gianni Pozzi

**Affiliations:** Laboratory of Molecular Microbiology and Biotechnology, Department of Medical Biotechnologies, University of Siena, 53100 Siena, Italy; Université Paris-Saclay, INRAE, AgroParisTech, Micalis, Jouy-en-Josas, France

**Keywords:** temperate phage, lysogenisation, *Streptococcus pyogenes*, *Streptococcus pneumoniae*, ϕ1207.3, mitomycin C, Transmission Electron Microscopy, qPCR

## Abstract

*Streptococcus pyogenes* prophage ϕ1207.3 (formerly Tn*1207.3*) carries the *mef*(A)-*msr*(D) efflux resistance genes, responsible for type M macrolide resistance. To investigate if ϕ1207.3 is a functional bacteriophage, we transferred the element from the original *S. pyogenes* host in a prophage-free and competence-deficient *S. pneumoniae* strain. Pneumococcal cultures of the ϕ1207.3-carrying lysogen were treated with mitomycin C to assess if ϕ1207.3 enters the lytic cycle. Mitomycin C induced a limited phage burst and a growth impairment resulting in early entrance in the stationary phase. To determine if ϕ1207.3 is able to produce mature phage particles we prepared concentrated supernatants recovered from a mitomycin C induced pneumococcal culture by sequential centrifugation and ultracentrifugation steps. Negative staining Transmission Electron Microscopy (TEM) of supernatants revealed the presence of phage particles with an icosahedral, electron dense capsid and a long, non-contractile tail, typical of a siphovirus. Quantification of ϕ1207.3 was performed by qPCR and semi-quantitatively by TEM. PCR quantified 3.34 × 10^4^ and 6.06 × 10^4^ excised forms of phage genome per ml of supernatant obtained from the untreated and mitomycin C treated cultures, respectively. By TEM, we estimated 3.02 × 10^3^ and 7.68 × 10^3^ phage particles per ml of supernatant. The phage preparations of ϕ1207.3 infected and lysogenised pneumococcal recipient strains at a frequency of 7.5 × 10^−6^ lysogens/recipient, but did not show sufficient lytic activity to form plaques. Phage lysogenisation efficiently occurred after 30 minutes of contact of the phages with the recipient cells and required a minimum of 10^3^ phage particles.

**Importance:** Bacteriophages play an important role in bacterial physiology and genome evolution. The widespread use of genome sequencing revealed that bacterial genomes can contain several different integrated temperate bacteriophages, which can constitute up to 20% of the genome. Most of these bacteriophages are only predicted *in silico* and never shown to be functional. In fact, it is often difficult to induce the lytic cycle of temperate bacteriophages. In this work, we show that ϕ1207.3, a peculiar bacteriophage originally from *Streptococcus pyogenes*, which can lysogenise different Streptococci and carries the macrolide resistance *mef*(A)-*msr*(D) gene pair, is capable of producing mature virions, but only at a low level, while not being able to produce plaques. This temperate phage is probably a partially functional phage, which seems to have lost lytic characteristics to specialize into lysogenisation. While we are not used to conceive phages separately from lysis, this behavior could actually be more frequent than expected.

## Introduction

Mobile genetic elements (MGEs) are segments of DNA that encode enzymes and proteins mediating their movement. All the MGEs are globally referred to as the mobilome (Siefert 2009), and represent a primary source of diversity for prokaryotes, since they can have profound effects on bacterial genome evolution by different means, such as by broadening the gene repertoire or by disrupting existing genes (Burrus and Waldor 2004; Johnson and Grossman 2015). These elements are also involved in the capture and spread of antimicrobial resistance genes. The MGEs usually implicated in the dissemination of antibiotic resistance genes are plasmids and Integrative Conjugative Elements (ICEs)(Botelho and Schulenburg 2021). Bacteriophages are the most abundant entities on the biosphere, with an estimated total number of virus-like particles on Earth close to 10^31^ (Hendrix et al. 1999; Clokie et al. 2011; Bar-On et al. 2018; Mushegian 2020). Nonetheless, phages are rarely found to carry antibiotic resistance genes (Enault et al. 2017) and their transfer rarely occurs between different bacterial species. We previously described a 52,491-bp genetic element integrated into the chromosome of the *Streptococcus pyogenes* clinical strain 2812A (Santagati et al. 2003; Pozzi et al. 2004). Since the element was transferred to other streptococcal species with a transfer mechanism resembling conjugation, it was named Tn*1207.3*. DNA sequence analysis showed that, beside the *mef*(A)-*msr*(D) macrolide efflux gene pair (Iannelli et al. 2014, 2018; Fox et al. 2021), the element contains phage genes and thus was renamed Φ1207.3. In *S. pyogenes* clinical isolates, other *mef*(A)-*msr*(D)-carrying prophages were found, including Φ10394.4, Φ29862, Φ29961, Φ29854, Φm46.1 and its variant VP_00501.1 (Banks et al. 2003; Brenciani et al. 2010; Di Luca et al. 2010; Vitali et al. 2016; Southon et al. 2020). To determine whether a prophage is fully functional, several properties of the element need to be checked. It should first be able to excise its genome, then to produce mature virions, and finally to infect new hosts and make plaques, or at least, lysogenise its host. To investigate if ϕ1207.3 is a fully functional bacteriophage, we transferred it in a *S. pneumoniae* prophage-free competence-deficient laboratory strain where we demonstrated that ϕ1207.3 is a bacteriophage able to excise, produce mature phage particles with a siphovirus morphology, and lysogenise a sensitive recipient.

## Materials and methods

### Bacterial strains and culture conditions

Strains used in this work and their relevant properties are reported in Table 1. Streptococcal strains were grown in Tryptic Soy Broth (TSB, BD) at 37°C. Starter cultures were sampled at an optical density at 590 nm (OD_590_) ranging from 0.2 to 0.3 and stored in 10% glycerol at −80°C. Solid media were obtained by supplementing TSB with 1.5% agar (BD) and 3% defibrinated horse blood (Liofilchem). When required, both liquid and solid media were supplemented with antibiotics at the following concentrations: 0.5 μg/ml erythromycin, 10 μg/ml novobiocin, 500 μg/ml streptomycin, 3 μg/ml chloramphenicol, 500 μg/ml kanamycin (Iannelli and Pozzi 2004).

**Table 1.** Bacterial strains and relevant properties.

| Strain | Properties | References |
| --- | --- | --- |
| 2812A | *Streptococcus pyogenes*, Italian clinical strain, serotype M1, carrying ϕ1207.3, Em<sup>R</sup> | (Santagati et al. 2003) |
| Rx1 | *Streptococcus pneumoniae*, rough highly transformable laboratory strain, *comC1-comD1*, Hex<sup>-</sup> | (Cuppone et al. 2021) |
| FP10 | Rx1 derivative, competence deficient (Δ*comC1*), *str-41*, Cm<sup>R</sup>, Sm<sup>R</sup> | (Iannelli et al. 2005, 2021; Santoro et al. 2010) |
| FP11 | Rx1 derivative, competence deficient (Δ*comC1*), *nov-1*, Cm<sup>R</sup>, Nov<sup>R</sup> | (Iannelli et al. 2005, 2021; Santoro et al. 2010) |
| FR1 | FP10 derivative carrying ϕ1207.3, Cm<sup>R</sup>, Sm<sup>R</sup>, Em<sup>R</sup> | This study |

Em, erythromycin; Cm, chloramphenicol; Sm, streptomycin; *comC1-comD1* encode the competence stimulating peptide and its receptor; Hex is the DNA mismatch repair system; *str-41* allele confers streptomycin resistance; *nov-1* allele confers novobiocin resistance.

### Transfer and lysogenisation assays

Transfer of bacteriophage ϕ1207.3 from *Streptococcus pyogenes* to *Streptococcus pneumoniae* was obtained through plate mating experiments (Iannelli et al. 2021). Briefly, donor and recipient cells were grown separately at 37°C in TSB in the presence of the appropriate antibiotics. Upon reaching the end of the exponential phase (OD_590_= 0.8), cells were mixed at a donor-recipient 1:10 ratio (i.e., 0.1 ml and 0.9 ml), centrifuged at 3,000 × *g* for 15 minutes, the pellet was resuspended in 0.1 ml of TSB and plated on TSA plates supplemented with 5% blood. Plates were incubated at 37°C in the presence of 5% CO_2_ up to 4 h, cells were then recovered with a cotton swab and resuspended in 1 ml of TSB supplemented with 10% glycerol and stored at −70°C. To select for recombinants, the mixture was then plated following a multilayer plating procedure, as previously described (Iannelli and Pozzi 2004; Iannelli et al. 2021). Recombinant phenotypes were confirmed by genetic analysis on plates containing the appropriate antibiotics and genotypes by PCR analysis (Santoro et al. 2010). To lysogenise pneumococcal recipients, a procedure similar to transfer assay was applied using phage preparation in place of the bacterial donor cells. For this, 0.9 ml of recipient cell culture (about 10^9^ CFUs) were centrifuged, the supernatant was discarded, and the pellet was resuspended directly in 0.1 ml of the phage preparation (see below), the mixture was incubated for 15 min at 37°C before plating on TSA plates supplemented with 5% blood, incubating for 30 minutes at 37°C, and recovering cells with a cotton swab. Selection of bacterial lysogens was performed by multilayer plating in the presence of 1 μg/ml erythromycin. The number of phage particles used for the lysogenisation assay was quantified by qPCR. To determine the minimum number of phage particles required for lysogenisation, the phage preparations were ten-fold serially diluted.

### Mitomycin C induction and phage preparations

Frozen starter pneumococcal culture was diluted 20-fold in 600 ml of TSB and grown at 37°C until reaching the early exponential phase (OD_590_ = 0.05-0.1), then the culture was split in two aliquots, of which one was treated with 100 ng/ml mitomycin C (Sigma Aldrich)(Bernheimer 1979; Romero et al. 2009). After 2 hours of incubation at 37°C, EDTA was added at a 10 mM final concentration and the culture was centrifuged two times at 5,000 × *g* for 40 minutes at 4°C in 50-ml tubes, to eliminate bacterial cells and cellular debris. The recovered supernatant of both induced and uninduced cultures was transferred in 6 polyallomer (36 ml each) centrifuge tubes (25×89 mm, Beckman Coulter) to which 1 ml of Protease Inhibitor Cocktail (Sigma Aldrich, P8465) was added before ultracentrifugation at 20,000 × *g* for 2 hours at 10°C in an Optima L-90K Ultracentrifuge with the SW32Ti rotor (Beckman Coulter). Supernatants were carefully discarded by aspiration and the 6 phage pellets were resuspended with 0.1 ml of TM buffer (50 mM Tris-HCl, 10 mM MgSO_4_) (Bernheimer 1979). A final volume of 0.7 ml was obtained combining the 6 suspensions. The fold concentration of the enriched phage preparation was calculated as follows: 216 ml of supernatant / 0.7 ml of suspension = 308.57x.

### Transmission Electron Microscopy

Phage preparations were observed by Transmission Electron Microscopy (TEM) after negative staining by placing 3 μl of phage preparation onto a glow-discharged Formvar-coated 300 mesh copper grid. The phage preparation was allowed to adsorb for 2 min, excess liquid was removed with filter paper, then a drop of 2% uranyl acetate was adsorbed on the grid and blotted dry. Samples were observed in a TECNAI G^2^ transmission electron microscope operated at 100 kV with magnification of 60,000×. Measurements of head diameter, tail length were made to verify the homogeneity of the phage preparation. About 10 representative meshes were observed for each preparation, and 200 fields were observed in each mesh, covering the entire mesh. The number of phage particles for each grid was then calculated by multiplying the average number of phages in a mesh by the number of meshes in a grid (300). The number of phage particles per ml of phage preparation was then calculated assuming all phages of the 3 μL drop had adsorbed to the grid, and used to quantify the theoretical number of phage particles per ml of bacterial culture supernatant.

### Bioinformatic analysis

The VIRFAM webserver (http://biodev.cea.fr/virfam/) for remote homology detection of viral family proteins (Lopes et al. 2014) was used to analyze the structural proteins of ϕ1207.3 and assess the family of the head-tail-neck module. Blast search was conducted in public protein databases and in Pfam protein family database using default parameters and only alignment with E-values below 0.001 were considered.

### Quantitative PCR

Quantitative Real-time PCR assays were performed with the KAPA SYBR FAST qPCR Master Mix (2X) Kit (Kapa Biosystems) on a LightCycler 1.5 (Roche) instrument, following the described procedure (Santoro et al. 2018, 2021). The reaction mix contained, in a final volume of 20 μl, 1X KAPA SYBR FAST qPCR reactive, each primer at a 200 μM final concentration. The following templates were used: i) the FR1 liquid culture, ii) the phage preparations from FR1 untreated culture, iii) the phage preparations from FR1 mitomycin C treated culture. Templates were incubated for 5 minutes at 90°C to allow bacterial cells and phage particles lysis and then 1 μl was added to the PCR reaction mix. The thermal cycling profile was as follows: 1 cycle of initial denaturation at 95°C for 3 minutes, 40 cycles of repeated denaturation 0 sec at 95°C, annealing 20 sec at 50°C, extension 10 sec at 72°C. The temperature transition rate was 20°C/s in the denaturation and annealing steps and 5°C/s in the polymerisation step. Melting curve was integrated at the end of the run by increasing temperature from 40°C to 95°C with a ramping of 0.05°C/s and acquiring fluorescence continuously. The primer pairs used were: (i) IF264/IF162, divergent primers, directed at the ends of the prophage genome, amplifying a fragment of 227 bp and used to quantify the excised phage; (ii) IF285/IF286, amplifying a 260 bp fragment, used for quantifying *mef(*A) gene; (iii) IF138/IF139, which amplified a 171 bp fragment of chromosomal *gyrB* gene, used for standardization of the results (Table S1). Serial dilutions of chromosomal DNA with known concentration were used to build a standard curve for the *gyrB* gene, by plotting the threshold cycle against the number of chromosome copies. This standard curve was recalled in the instrument software to standardize the number of *mef*(A) gene copies and excised phage genomes. Analysis of the melting curves was performed to discern desired amplification products from primer dimer products. The number of excised phage genomes copies per ml of phage preparation was converted to number of excised phage genomes copies per ml of bacterial supernatant. Fold enrichment was calculated as the ratio between the value obtained in the phage preparation and in the bacterial culture.

### Double Agar Overlay Plaque Assay

In the plaque assay, 0.05 ml of the phage preparation (about 1.18 × 10^5^ phage particles, as quantified by TEM) were added to 0.1 ml of pneumococcal culture, sampled in exponential phase (OD_590_ = 0.2-0.3), of the indicator strain FP10. The phage-bacteria mixture was added to 3 ml of the top agar (TSB/0.4% Seakem LE agarose), mixed gently, and poured into a 90-mm petri dish containing 25 ml of TSA supplemented with 10 mM CaCl_2_. The plates were dried for 10 min at room temperature and incubated at 37°C overnight in a 5% CO_2_ enriched atmosphere (Kropinski et al. 2009).

## Results and discussion

### Transfer of ϕ1207.3 to Streptococcus pneumoniae

To investigate if ϕ1207.3 is a functional bacteriophage, the genetic element was transferred from the original clinical *Streptococcus pyogenes* host in a prophage-free, competence-deficient pneumococcal strain. The resulting pneumococcal FR1 lysogen strain was obtained with a mating experiment where *S. pyogenes* 2812A was the ϕ1207.3 donor and *S. pneumoniae* FP10 was the recipient (Table 1). Transfer of ϕ1207.3 occurred at a frequency of 3.8 × 10^−5^ ± 7.6 × 10^−6^ lysogens per donor, as already observed (Iannelli et al. 2021).

### φ1207.3 induction by mitomycin C

To assess if the ϕ1207.3 enters the lytic cycle, liquid cultures of the ϕ1207.3-carrying strain FR1 were induced with mitomycin C at sub-inhibitory concentration. Mitomycin C induced a limited phage burst and a growth impairment resulting in early entrance in the stationary phase (Figure 1). FR1 generation time was 51 minutes in the absence, and 70 minutes in the presence of the mitomycin C stimulus. In the untreated cultures, the stationary phase was reached after 205 minutes, corresponding to 4 generations, whereas in the treated culture it was reached after 121 minutes, corresponding to 1.6 generations. FP10 generation time was 54 minutes in the absence, and 59 minutes in the presence of mitomycin C. In both the untreated and mitomycin C treated cultures, the stationary phase was reached after 180 minutes, corresponding, respectively, to 3 and 3.3 generations. Hence, although bacteriophage ϕ1207.3 does not seem to propagate efficiently by lytic cycle on plates, as it does not form lysis plaques, mitomycin C caused a significant, phage-specific growth impairment, as the treated culture reached the stationary phase 2.4 generations before the untreated culture. Almost no impairment of growth was observed in the isogenic progenitor control strain FP10 after mitomycin C treatment at a sub-inhibitory concentration (Tomasz 1995).

**Figure 1.**
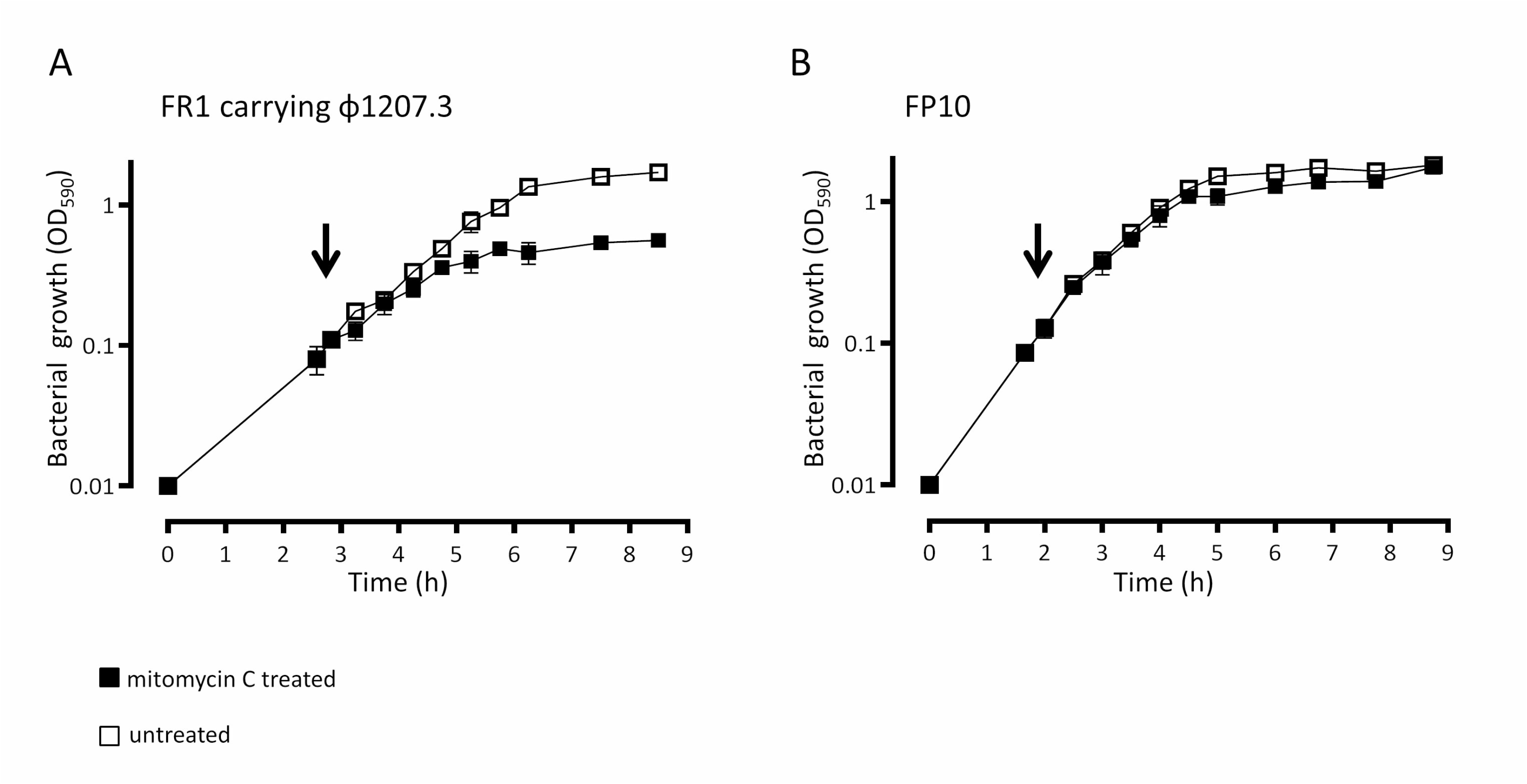
ϕ1207.3 induction by mitomycin C. To determine if ϕ1207.3 has a lytic phenotype, mitomycin C was added, at a sub-inhibitory concentration (100 ng/ml), to liquid bacterial cultures in early exponential phase. A) In the ϕ1207.3-carrying strain FR1, mitomycin C causes a limited phage burst and a growth impairment resulting in early entrance in the stationary phase (2.4 generations before of the untreated culture). B) In the FP10 parental strain mitomycin C produces only a slight growth impairment. The time point of mitomycin C addition is indicated by an arrow. Results are reported as means and standard deviations resulting from at least three independent experiments.

### Detection of φ1207.3 phage particles

In order to determine if ϕ1207.3 is able to produce mature phage particles, we prepared supernatants from the ϕ1207.3-carrying strain FR1. The supernatant recovered from a culture treated or not with mitomycin C was concentrated by sequential centrifugation and ultracentrifugation steps, without filtering to minimize particle breakage. Observation of the ultracentrifuged supernatants at the TEM showed the presence of phage particles with an icosahedral, electron dense capsid and a long, non-contractile tail, with tail fibers visible in some images. The capsid was found to be 62 nm in diameter, while the tail was 175 nm (± 1 nm) in length and 8 nm in diameter. The morphology observed is consistent with a siphovirus virion (Figure 2). We specifically avoided a filtering step since it could significantly decrease phage recovery (Castro-Mejía et al. 2015), moreover we ultracentrifuged samples at a relatively low speed (20,000 × *g*) to preserve the integrity of viral particles (Bourdin et al. 2014). We conclude that ϕ1207.3 is able to produce mature phage particles, suggesting it is not a defective prophage.

**Figure 2.**
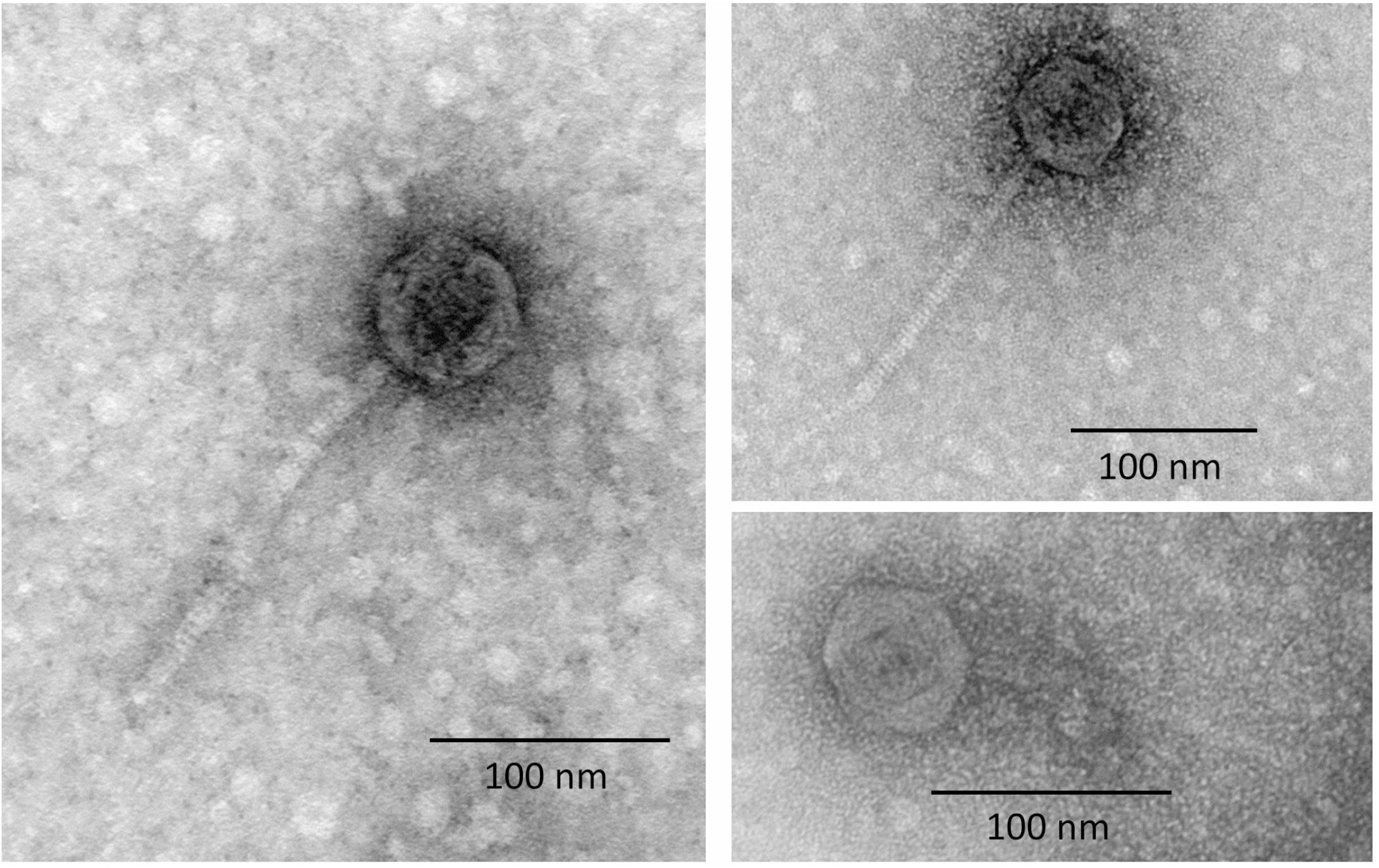
Transmission Electron Microscopy of the Φ1207.3 phage particles. TEM observation after negative staining of phage preparations shows the presence of phage particles with a morphology consistent with a siphovirus. The capsid shows an icosahedral symmetry and had a diameter of 62 nm while the tail was non-contractile and 175 nm (± 1 nm) long. Scale is reported for each panel.

### Sequence analysis of φ1207.3 predicted proteins

To integrate the structural data from TEM, the head-neck-tail module search of Virfam was used to predict the ϕ1207.3 phage morphology. Virfam analysis assigned ϕ1207.3 to Siphoviridae with a cluster 2 type 1 neck, a family of phages that infect *Firmicutes* bacteria. Virfam analysis and Blast homologies search, conducted in public protein databases and in Pfam protein family database, predicted Orf38 to Orf47, Orf50, Orf52 and Orf53 as putative structural proteins involved in the assembly of a complete phage particle (Table S2). In addition, and as already observed, a predicted lysis module was found, composed by *orf53* and *orf54* coding for predicted holin and amidase proteins, respectively (Table S2) (Iannelli et al 2014).

### Quantification of φ1207.3 in enriched phage preparations

Quantification of ϕ1207.3 was performed by qPCR and with a semi-quantitative approach based on the TEM imaging, in phage preparations from both untreated and mitomycin C treated FR1 cultures. Phage preparations, obtained after sequential centrifugation steps, were concentrated 308.57-fold compared to the original culture supernatant. We used qPCR to quantify the copy number of (i) the joints between the ends of ϕ1207.3 prophage sequence, i.e., excised form of ϕ1207.3 genome, (ii) the macrolide resistance gene *mef*(A) gene carried by ϕ1207.3, and (iii) the chromosomal reference gene *gyrB*, in both phage preparations and bacterial cultures (Table 2). We estimated 1.03 × 10^7^ excised forms of ϕ1207.3 genome per ml in the concentrated phage preparations obtained from the untreated cultures and 1.87 × 10^7^ excised forms per ml in the phage preparations from the mitomycin C treated cultures. These values corresponded, respectively, to 3.34 × 10^4^ and 6.06 × 10^4^ excised forms per ml of the original culture supernatant. Within both phage preparations, from untreated and mitomycin C treated cultures, a 20- to 25-fold enrichment of phage genes compared to the bacterial *gyrB* gene was observed, proving that most of the phage signal does not come from contaminating bacterial DNA. Moreover, an enrichment in excised forms and *mef*(A) gene copy number was detected when compared to the original bacterial culture. The excised forms were enriched 38-fold in phage preparations obtained from the untreated cultures compared to the bacterial culture, while the enrichment was 70-fold in the phage preparations obtained from mitomycin C treated cultures. The *mef*(A) gene was enriched 7.8- and 38-fold, respectively. In contrast, a 10- and 5-fold decrease was observed for the chromosomal reference *gyrB* gene. To complement qPCR quantification, a semi-quantitative approach based on the TEM was implemented to quantify the number of ϕ1207.3 particles in the culture supernatant. We observed and counted phages in 30 meshes, from at least three independent experiments. We estimated about 9.33 × 10^5^ ± 2.31 × 10^5^ phage particles per ml in concentrated phage preparations obtained from untreated cultures and 2.37 × 10^6^ ± 9.96 × 10^5^ phage particles per ml in the concentrated phage preparations from mitomycin C treated cultures. These values corresponded, respectively, to 3.02 × 10^3^ and 7.68 × 10^3^ phage particles per ml of the original bacterial supernatant. The qPCR estimate was in general about 10 times higher, compared to the TEM estimate. The discrepancy is possibly explained by the presence of free phage DNA in the preparation or, less likely, by the multiple genome copies packed in the phage head (Zhang et al. 2011). However, in both cases, the fold increase upon mitomycin C treatment was similar. We conclude that mitomycin C, while having a clear and specific effect on the lysogen culture suggestive of prophage induction, resulted in a moderate production of viral particles (about a two fold increase, compared to uninduced cultures), suggesting that the phage lytic cycle was somewhat impeded.

**Table 2.**
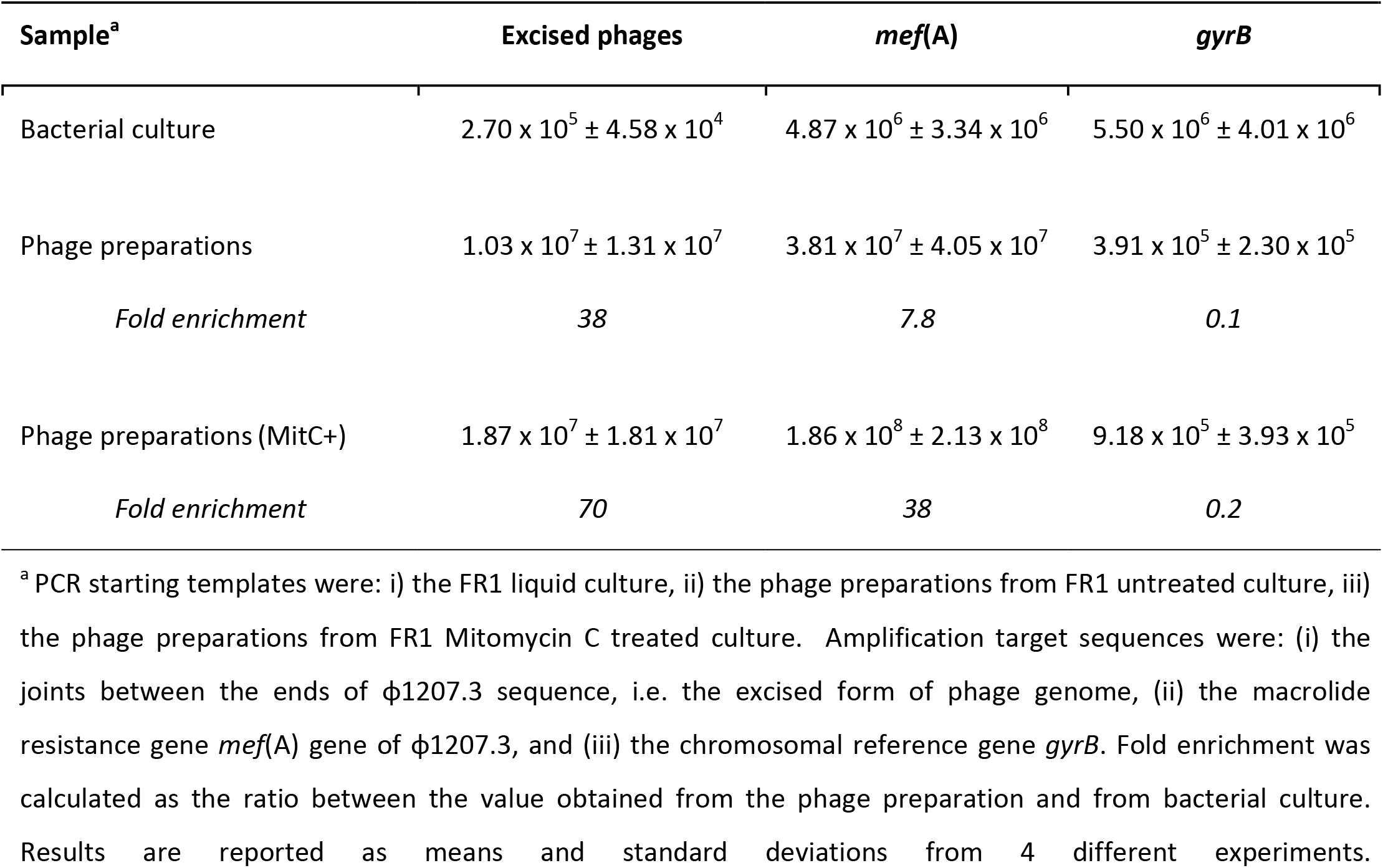
PCR quantification of Φ1207.3 phage genome.

### Absence of plaque formation

To further assess whether ϕ1207.3 performs lytic cycles, phage preparations obtained from a mitomycin C treated FR1 bacterial culture were used in plaque assay experiments. No plaques were observed with the indicator *S. pneumoniae* strain FP11. We conclude that, in our experimental conditions, a moderate cell lysis is taking place in liquid culture upon mitomycin C treatment (Figure 1), but no plaque formation is detectable on plates of strain FP11. This is suggestive of a phage having a low burst size, or a preference for growth in liquid cultures.

### Lysogenisation with φ1207.3

Finally, we tested if ϕ1207.3 can infect and lysogenise *S. pneumoniae* FP11. For this purpose, we used concentrated phage preparations obtained from mitomycin C treated culture, placed them into contact with the novobiocin resistant and competence deficient *S. pneumoniae* FP11 strain. Lysogens were obtained at an average frequency of 7.5 × 10^−6^ ± 5.7 × 10^−6^ per recipient, indicating that ϕ1207.3 phage particles are able to lysogenise *S. pneumoniae*. The minimum number of phages required to obtain a detectable lysogenisation was 10^3^, whereas no lysogens could be detected (<2.8×10^−8^ per recipient) using lower phage numbers (Figure 3). Since we did not include a filtering step, our phage preparations also contained a residual number of bacteria (about 5 × 10^4^ CFU/ml) which were counter-selected using novobiocin as a selection marker. As a further control, to rule out the contribution of whole bacterial cells to phage lysogenization (i.e. a conjugation-mediated transfer), we performed transfer experiments using ten-fold serial dilutions of an FR1 bacterial culture. When less than 6 × 10^6^ CFUs of donor bacteria were used, no phage transfer was observed (< 8.3 × 10^−9^ lysogens per recipient). Finally, we assessed the kinetics of phage lysogenization in a time course experiment, varying the lysogenisation incubation time from 30 to 240 minutes (Figure 4). The number of lysogens was stable over time, indicating that lysogenisation efficiently occurs after 30 minutes from the contact of the phages with the recipient cells. On the other hand, the lysogenisation frequency decreased over time as a consequence of increasing recipient CFUs.

**Figure 3.**
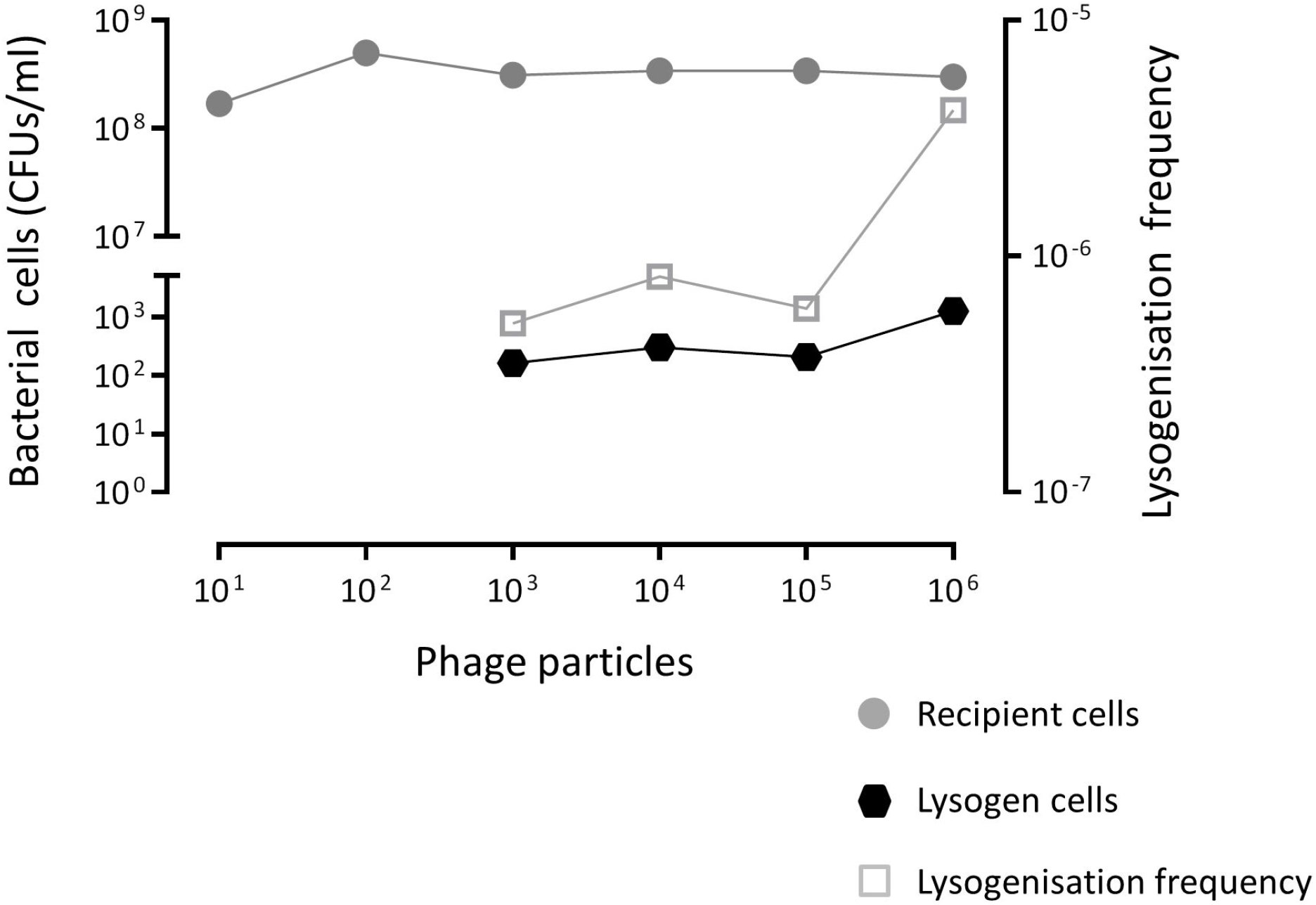
Lysogenisation by ϕ1207.3. Ten-fold serially diluted phage preparations obtained from mitomycin C treated FR1 culture were used to lysogenise the novobiocin resistant and competence deficient *S. pneumoniae* FP11. The minimum number of phages required to obtain a detectable lysogenisation was 10^3^, whereas no lysogens could be detected (<2.8×10^−8^ lysogens per recipient) using lower phage numbers. The lysogenisation frequency was calculated as number of lysogens per recipient cell.

**Figure 4.**
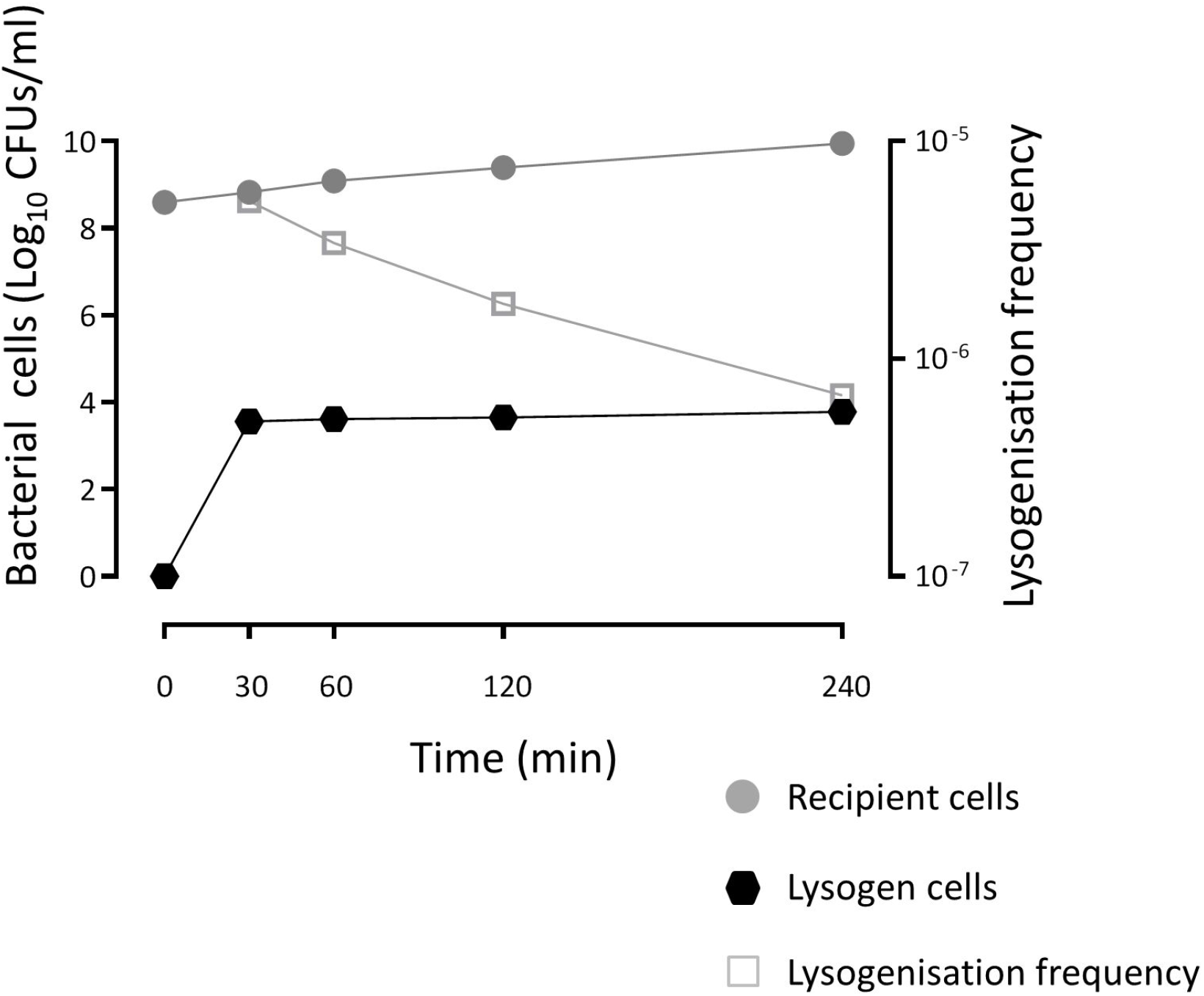
Kinetics of lysogenisation by ϕ1207.3. A time-course lysogenisation experiment shows that the number of recombinant lysogens is stable over time, indicating that lysogenisation efficiently occurs within 30 minutes after the contact of the phages with the recipient cells. The lysogenisation frequency was calculated as the number of lysogens per recipient cell.

## Conclusions

Prophages play an important role in shaping the *S. pyogenes* genome affecting the physiology and pathogenesis of this organism. Prophages are rarely found to directly encode antimicrobial resistance genes and their transfer seldom occurs between different bacterial species (Giovanetti et al. 2015; Enault et al. 2017; Abouelfetouh et al. 2022). In *S. pyogenes*, the macrolide efflux genes *mef*(A)-*msr*(D) were found associated to several prophages, including ϕ1207.3, Φ10394.4, Φ29862, Φ29961, Φ29854, Φm46.1 and its variant VP_00501.1 (Banks et al. 2003; Brenciani et al. 2010; Vitali et al. 2016; Southon et al. 2020). We found ϕ1207.3 in the M1 serotype *S. pyogenes* clinical strain 2812A exhibiting the M phenotype of resistance to macrolides (Santagati, 2003). Even if *S. pyogenes* strains devoid of phages were reported (Beres et al. 2016), most *S. pyogenes* genomes contain at least 2 prophages, with a maximum of 8 found in the genome of MGAS10394 (Banks et al. 2004). The genome of *S. pyogenes* 2812A contains other 2 prophages homologous to ϕ315.3 and to ϕ5005.3 (our unpublished data). Since it is difficult to study the biology of a single prophage in the *S. pyogenes* original host due to its polylysogeny, ϕ1207.3 was transferred to a prophage-free competence-deficient *S. pneumoniae* laboratory strain. We demonstrated that ϕ1207.3 is able to excise its genome, to produce mature phage particles in *S. pneumoniae*, with a morphology consistent with that of a siphovirus, and to lysogenise sensitive recipients. In the *S. pneumoniae* lysogen hosting ϕ1207.3, mitomycin C treatment does not induce a full lysis, but only a limited growth arrest, and phage particle production is only two-fold above the uninduced strain level. Lysogenisation by ϕ1207.3 was obtained with concentrated phage preparations containing at least 10^3^ phage particles, while it could not be obtained using the non-concentrated culture supernatant (Santagati et al. 2003).

Furthermore, lysogenisation could only be detected when bacterial cells and phage particles were in close contact on an agar plate, suggesting that biofilm formation may be important for the transfer of this phage (Madsen et al. 2012; Lécuyer et al. 2018). In conclusion, we demonstrated that ϕ1207.3 is a partially functional phage with peculiar characteristics i.e. (i) the association with antibiotic resistance determinants, (ii) the ability to lysogenise several *S. pyogenes* strains with different genetic backgrounds (our unpublished data) and other streptococcal species such as *S. gordonii* and *S. pneumoniae* (Santagati et al. 2003), (iii) the impairment in producing plaques on plate. Further investigations, e.g. in the original *S. pyogenes* host devoid of prophages (Euler et al. 2016), are required to fully elucidate the mechanism underlying the activities of this unusual bacteriophage capable of lysogenisation while inefficient in its lytic cycle, as if upon infection, the choice between lysis and lysogeny was tilted heavily towards lysogeny.

## Acknowledgments

We thank Eugenio Paccagnini and Pietro Lupetti (Department of Life Sciences, University of Siena) for their help with Transmission Electron Microscopy.

This work was supported in part from the Italian Ministry of Education, University and Research (MIUR-Italy) under grant number 20177J5Y3P (call “Progetti di Ricerca di Rilevante Interesse Nazionale – Bando 2017”), and in part from the European Union Seventh Framework Programme (FP7/2007-2013) under grant agreement 241446 (project ANTIRESDEV).

